# Patient-specific therapeutic benefit of MuSK agonist antibody ARGX-119 in MuSK myasthenia gravis passive transfer models

**DOI:** 10.1101/2024.08.01.606156

**Authors:** Jamie L. Lim, Stine Marie Jensen, Jaap J. Plomp, Bernhardt Vankerckhoven, Christa Kneip, Rani Coppejans, Christophe Steyaert, Kathleen Moens, Lieselot De Clercq, Martijn R. Tannemaat, Peter Ulrichts, Karen Silence, Silvère M. van der Maarel, Dana L.E. Vergoossen, Roeland Vanhauwaert, Jan. J. Verschuuren, Maartje G. Huijbers

## Abstract

Muscle-specific kinase (MuSK) orchestrates establishment and maintenance of neuromuscular synapses, which enable muscle contraction. Autoantibodies targeting MuSK cause myasthenia gravis (MG), a disease characterized by fatigable skeletal muscle weakness which requires chronic immunosuppressive treatment and ventilatory support at some point in ∼30% of patients. MuSK autoantibodies are predominantly IgG4 and are bispecific, functionally monovalent antibodies due to Fab-arm exchange. Through monovalent binding, MuSK IgG4 autoantibodies act as antagonists on the MuSK signalling pathway, impairing neuromuscular synaptic function. In contrast, bivalent MuSK antibodies act as agonists of the MuSK signalling pathway. Since symptoms in MuSK MG are largely caused by antagonistic monovalent MuSK antibodies, we hypothesized that a bivalent MuSK agonist could rescue MuSK MG, bypassing the need for generalized immunosuppression. In this study, we investigated whether an agonist antibody targeting the Frizzled-like domain of MuSK, ARGX-119, can ameliorate disease in MuSK MG models induced by passive transfer of polyclonal IgG4 from unrelated patients. For each patient material we first established the minimal dose for a progressive MG phenotype based on muscle function tests. ARGX-119 significantly improved survival and muscle weakness in a mouse model induced by one patient material, but not by three others. Mechanistically, this patient-specific efficacy could not be explained by autoantibody epitope specificity, titer or competition for ARGX-119 binding, but rather correlated to the presence of MuSK activating antibodies in some patients. We further provide evidence that an *in vitro* assay may predict which patients potentially benefit from ARGX-119 and that this treatment, when effective in MuSK MG mice, follows a bell-shaped dose-effect curve. These results provide first proof of concept of a MuSK agonist in a clinically relevant model for MuSK MG. We anticipate this to be a starting point for investigating the therapeutic benefit of ARGX-119 in MuSK MG and other neuromuscular diseases hallmarked by neuromuscular synaptic dysfunction.

**Graphical abstract:** 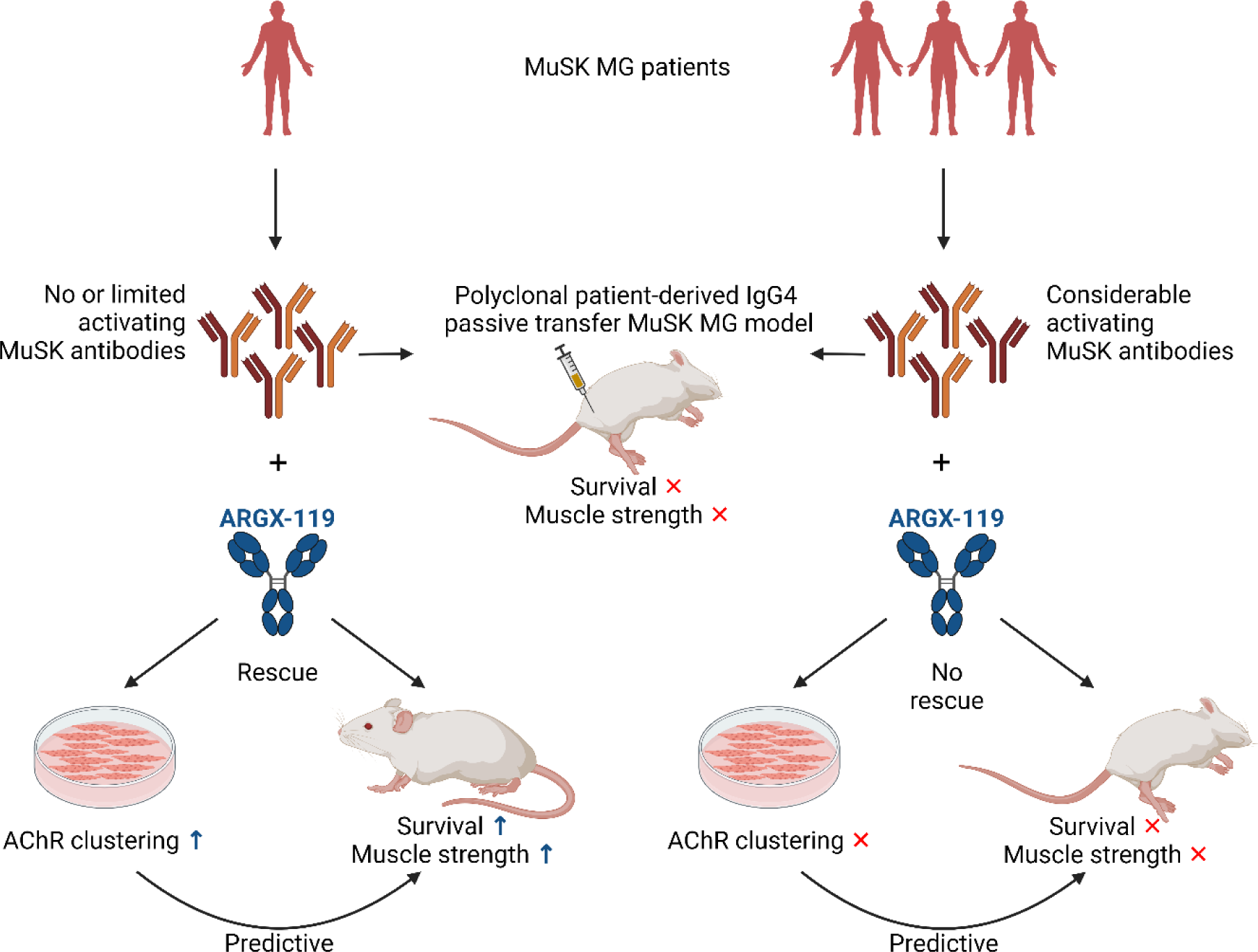

**Highlights:**

- MuSK agonist ARGX-119 can rescue MuSK MG in a patient-specific manner
- MuSK agonism follows a bell-shaped efficacy curve in this MuSK MG mouse model
- Variation in ARGX-119 efficacy between patient models is not explained by competition for binding on MuSK, but rather appears related to an agonistic fraction of patient antibodies
- An *in vitro* assay is potentially predictive for treatment efficacy of the MuSK agonist

## Introduction

Myasthenia gravis (MG) is one of the most prevalent neuromuscular diseases, yearly affecting 10-29 per million individuals.^1, 2^ Clinically, MG is characterized by asymmetric ocular muscle weakness and fluctuating and fatigable skeletal muscle weakness. The disease is caused by autoantibodies impairing neuromuscular synaptic communication by blocking the function of, or reducing the number of essential neuromuscular synaptic proteins, such as acetylcholine receptors (AChRs). In 2001, autoantibodies to muscle-specific kinase (MuSK) were shown to be associated with ∼5% of MG cases.^3, 4^

MuSK is a postsynaptic muscle membrane molecule that is a master regulator of neuromuscular synapse formation.^5–7^ It is furthermore needed to maintain these synapses throughout life. The MuSK signaling pathway, among others, induces AChR clustering, which is crucial for neuromuscular transmission, and thus enables muscle contraction.^8, 9^ MuSK, through clustering low density lipoprotein receptor-related protein 4 (Lrp4), furthermore instructs the motor nerve terminal to remain differentiated.^10^ It is therefore critical that active MuSK signaling is maintained continuously through Lrp4 and agrin binding.^11^ Perturbation of MuSK signaling e.g. through mutations in *MUSK*, or through autoantibodies binding to MuSK, causes impairment of muscle function in congenital myasthenic syndrome (CMS)^12^ or autoimmune MG, respectively.^3, 13–16^

The dominant epitope for autoantibodies causing MuSK MG is the N-terminal Ig-like domain 1 of MuSK, although more than half of the patients have additional antibodies to other domains.^17, 18^ The Ig-like domain 1 is critical for interaction with Lrp4.^19^ MuSK autoantibodies obstruct the interaction between MuSK and Lrp4 and thereby prevent the activation of the MuSK signaling cascade.^20–22^

Surprisingly, MuSK autoantibodies in MG are predominantly of the IgG4 subclass. IgG4 is considered an anti-inflammatory antibody type, as it has limited capacity to activate complement^23^ and stimulates inhibitory rather than activating Fcγ receptors on immune cells.^24^ In addition, IgG4 exchanges half molecules in the circulation in a stochastic process called Fab-arm exchange.^25^ The net effect of this is that ∼99% of IgG4 antibodies in circulation are effectively bispecific and engage in monovalent antigen binding. Indeed, MuSK autoantibodies were also shown to be functionally monovalent in patients.^26^ Notably, functionally monovalent MuSK antibodies, based on patient-derived autoantibody sequences, were found to be much more pathogenic than their bivalent parental equivalent.^27^ These differences can be explained by their opposing effects on MuSK signaling. Functionally monovalent MuSK antibodies block agrin and Lrp4-mediated MuSK activation and AChR clustering, whereas bivalent MuSK antibodies have agonistic potential on MuSK signaling and can independently induce AChR clustering.^27, 28^ Since symptoms in MuSK MG are largely caused by antagonistic monovalent Ig-like domain 1 MuSK antibodies, we hypothesized that a bivalent MuSK agonist could rescue MuSK MG. Here we investigate whether an engineered MuSK agonist antibody, ARGX-119, binding the Frizzled-like domain (Fz-domain) of MuSK, can rescue MuSK MG induced by polyclonal patient IgG4.

## Results

### Patient-specific rescue of MuSK MG in mice using ARGX-119

To establish mouse models to test ARGX-119 efficacy, we first determined the minimal dose per patient purified IgG4 for a maximal MG phenotype. We reasoned that this dosing strategy provides the largest therapeutic window for ARGX-119 as a minimal treatment effect should result in a change in phenotype. Potency for induction of myasthenic symptoms varied between patient IgG4s (patient 2 > patient 1 > patient 3 > patient 4), with the minimal dose ranging from 25 mg/kg to 60 mg/kg (Supplementary Figure 1), which appears to be (partly) due to differences in reactivity of the patient IgG4s as determined by MuSK ELISA (Supplementary Table 1). Therefore, the following ‘disease doses’ were used in the therapeutic passive transfer MuSK MG experiments: 40 mg/kg of patient 1, 25 mg/kg of patient 2, 50 mg/kg of patient 3 and 60 mg/kg of patient 4 material.

Passive transfer mouse MuSK MG models using daily injections of patient IgG(4) have a fast onset and can be considered severe, as they cause progressive synaptic disintegration and (subclinical) symptoms after one day.^14, 29^ Therefore, we reasoned that prophylactic treatment with ARGX-119 or Mota one day prior to the start of daily injections of IgG4 patient antibodies, followed by ARGX-119 or Mota every 3 days, would be a good starting treatment regimen (Figure 1A). Exposure to 2.5 mg/kg ARGX-119 demonstrated significant therapeutic efficacy versus 2.5 mg/kg isotype control Mota in the MuSK MG mouse model induced with IgG4 from patient 1. The ARGX-119 treated mice improved on all functional parameters, including survival and motor performance as measured by forepaw grip strength and hanging time (Figure 1B), despite continued patient antibody dosing. Progressive loss of body weight started from day 8 in Mota mice, while body weight stabilized above 90% in the ARGX-119 treated mice until end of the study (Supplementary Figure 2). As a result, five out of 6 (83%) ARGX-119 treated mice survived the experiment (did not reach the humane endpoint of ≥20% body weight loss), compared to one out of six (17%) in the Mota group. The surviving Mota treated mouse had an overall milder disease profile compared to the other mice in the group. The same therapeutic effect of ARGX-119 was observed for grip strength (∼30% increase in grip strength in ARGX-119 versus Mota treated mice) and hanging time (120-140 seconds increase in hanging time in ARGX-119 versus Mota treated mice). When the same dose of ARGX-119 was tested in the MuSK MG model induced by IgG4 from three other patients, no statistically significant therapeutic efficacy of ARGX-119 was observed (Figure 1B, Supplementary Figure 2). A trend for ARGX-119 efficacy was seen in all outcome parameters with IgG4 from patient 3, which had a less severe disease profile. In addition, ARGX-119 treated groups performed consistently slightly better on survival. These results, however, were not as evident as the therapeutic effect of ARGX-119 in the model using patient 1 IgG4.

**Figure 1:**
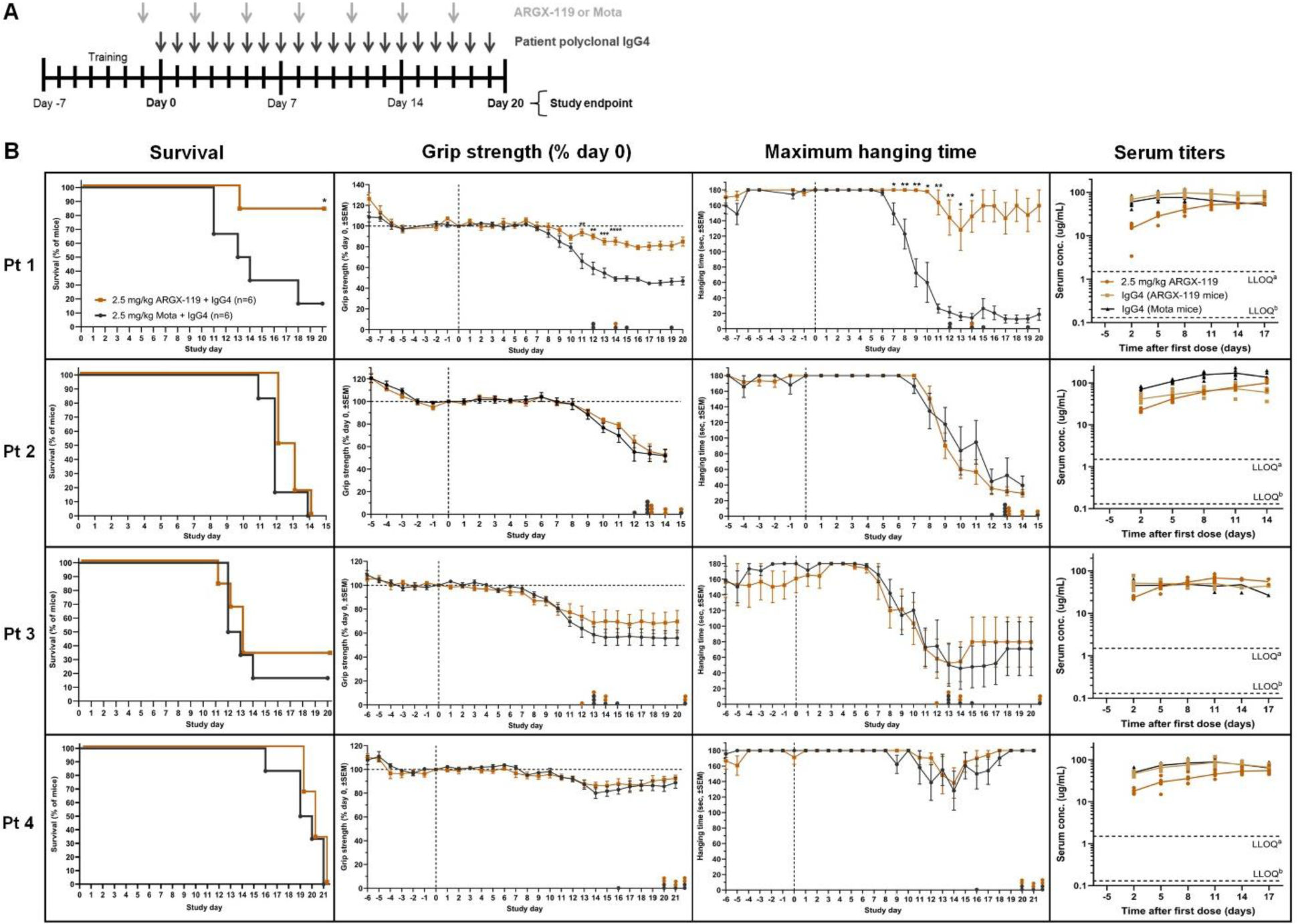
ARGX-119 rescues the MuSK MG phenotype induced with polyclonal patient IgG4 in a patient-specific manner. A) MuSK MG mouse model study design. B) *From left to right for patients IgG4 1-4*: percentage mouse survival over time, with reductions in survival representing sacrifice of mice due to humane endpoint reached; percentage grip strength normalized to day 0; hanging time (maximum 180 seconds, best of 3 attempts); and ARGX-119 and patient IgG4 serum concentrations (ug/mL) of mice treated with 2.5 mg/kg ARGX-119 (n=6) or isotype control Mota (n=6). Data points of percentage grip strength and hanging time represent grouped values of the mean ±SEM of mice. In case of missing data due to mouse sacrifice, the last value of each mouse was carried forward until the end of the experiment. Data points on the x-axis of outcome parameters represent mice with missing data after reaching humane endpoint or end of study. Statistical analyses were only performed on average data points representing three or more mice in each group. *P<0.05, **P<0.01, ***P<0.001, ****P<0.0001 for ARGX-119 vs Mota treated mice.

Pathogenic potential of the unrelated patient IgG4s varied significantly in these experiments, despite previous dose-finding experiments. This may be related to the inherent variation in mice exposed to the same patient antibody dose as well as the relatively low numbers of mice in the dose-finding experiments due to limiting amounts of available polyclonal patient IgG4. For patient 3 and 4, the lethal disease dose for some animals was not reached as some seemed to recover independent of the treatment. This is likely a reflection of using an IgG4 dose that’s just on the verge of being pathogenic, with some animals progressing to full blown myasthenia, while others are more resistant or capable of compensating the autoantibody attack, an experimental variable we’ve previously observed as well.^14, 30^ To exclude that a lack of efficacy of ARGX-119, or a reduced disease severity, in these animals was caused by altered exposure, we determined serum concentrations of ARGX-119 and patient IgG4 per model (Figure 1B, Supplementary Figure 3). In each model, only one or two mice had slightly lower serum patient IgG4 titers compared with the other mice, which may reflect individual differences in pharmacokinetics. However, exposure to ARGX-119 and patient antibodies was overall consistent throughout experiments and no differences were observed between treatment groups that could explain the differences in therapeutic efficacy or disease severity between models. In conclusion, ARGX-119 gave pronounced improvement of myasthenic muscle weakness in one out of four patient polyclonal IgG4-based MuSK MG mouse models.

### ARGX-119 has a dose-dependent, bell-shaped efficacy in the MuSK MG mouse model

Successful MuSK agonism is dependent on a (bivalent) antibody forcing dimerization of two MuSK molecules and activation of the kinase domains.^31–33^ Antibody-mediated agonism through dimerization, such as with ARGX-119, generally shows an optimum, as too high doses result in monovalent antibody binding and loss of efficacy (i.e. resulting in a bell-shaped dose-effect curve).^34^ Synaptic MuSK levels may be reduced in MuSK MG patients and mice,^29^ and this may result in sensitivity to over-dosing of a MuSK agonist due to monovalent binding. To determine optimal dosing of ARGX-119, we used polyclonal IgG4 from patient 1 to induce MG in mice as described above (Figure 1A) and exposed the animals to a dose range of ARGX-119. Lower (0.1 mg/kg) or higher (10 mg/kg) doses of ARGX-119 gave moderate improvement of the disease symptoms over time, including body weight loss and muscle strength/fatigability (% grip strength and hanging time), whereas 0.01 mg/kg and 20 mg/kg had little to no treatment effect (Figure 2A-C). In addition, these experiments revealed that ARGX-119 has an optimum efficacy at 0.25-2.5 mg/kg in this disease model (Figure 2D-F), supporting the experimental design used for the model described in Figure 1. The efficacious range for ARGX-119 is therefore estimated to span 2-3 orders of magnitude, with an optimum spanning between 1 and 2 orders of magnitude in this model. These data confirm the dose response curve of ARGX-119 in this MuSK MG mouse model to be bell-shaped.

**Figure 2:**
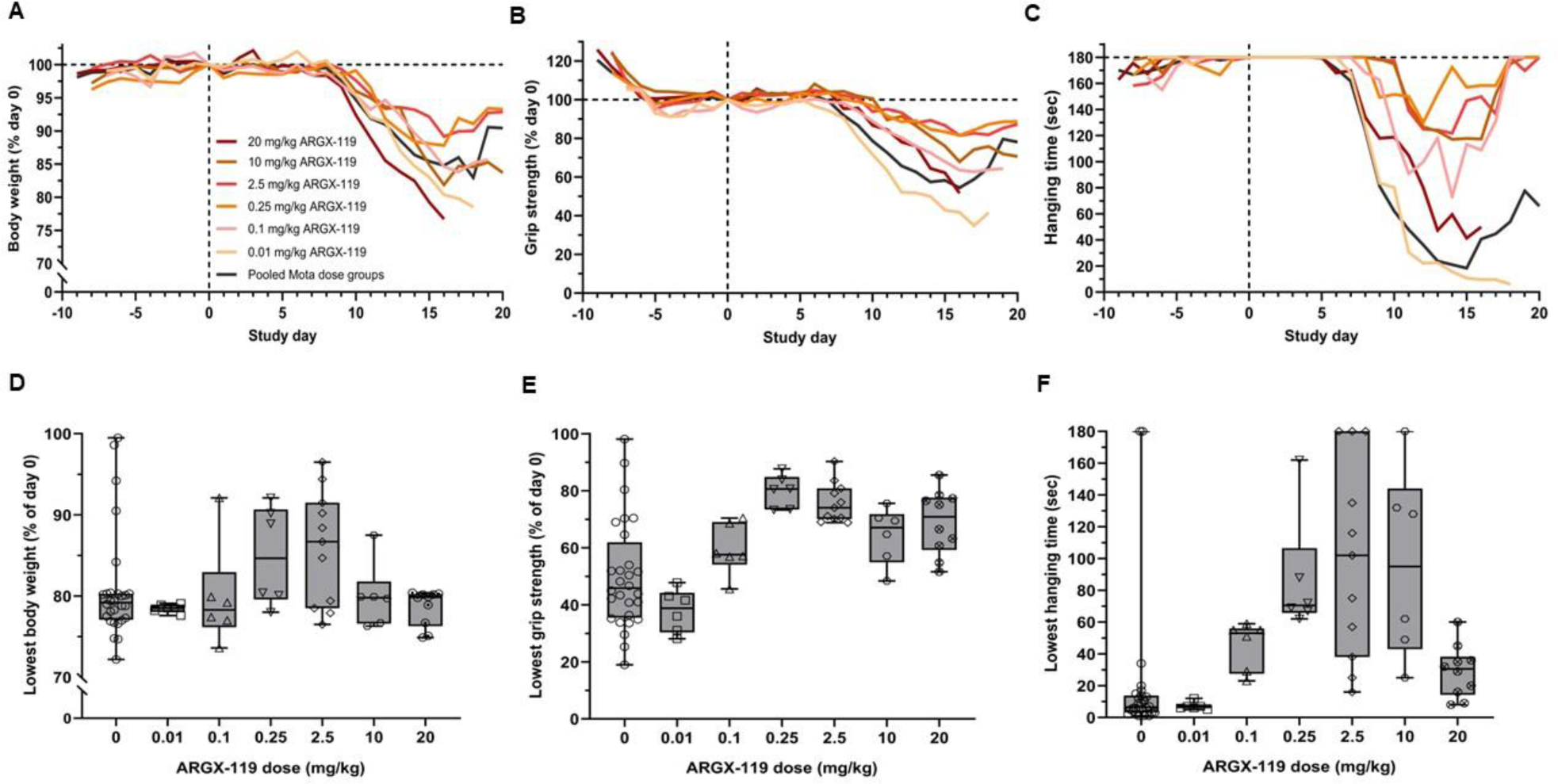
ARGX-119 shows a bell-shaped dose-effect relationship in the MuSK MG passive transfer model with Pt 1 IgG4. A) percentage body weight of day 0 over time, B) percentage grip strength of day 0 over time, C) hanging time (maximum 180 sec, best of three attempts) over time, D) lowest percentage body weight of day 0, E) lowest percentage grip strength of day 0, F) lowest hanging time (maximum 180 sec, best of three attempts) for 0 mg/kg ARGX-119 (pooled Mota treated mice, n=28), 0.01 mg/kg (n=6), 0.1 mg/kg (n=6), 0.25 mg/kg (n=6), 2.5 mg/kg (n=11), 10 mg/kg (n=6), 20 mg/kg (n=10). Horizontal line within each boxplot is the median, the ends of the box are the 25th and 75th percentiles and the whiskers show the lowest or highest data value still within 1.5-fold of the lower or upper interquartile ranges, respectively. Data points represent individual mice within the dose groups.

To exclude that we had missed the therapeutic optimum for ARGX-119 in our first experiments with the three non-responsive patient materials, we performed additional mouse experiments with these patient IgG4s using dosing of ARGX-119 at 0.25 mg/kg (Supplementary Figure 4). Data from these experiments demonstrated that similar to the 2.5 mg/kg dose, significant improvement on body weight, survival, grip strength and hanging time parameters was achieved versus the Mota control in the mouse MuSK MG model induced with patient 1 IgG4, but not with IgG4 from the other three patients, except for a trend of slight improvement in the model using IgG4 from patient 3. Thus, we determined the optimal dose and confirmed a bell-shaped dose-effect curve for MuSK agonism in one MuSK MG mouse model. This additional understanding of MuSK agonist dosing did not, however, bring out significant therapeutic efficacy in the MuSK MG models using patient materials 2-4.

### *In vitro* assessment of patient-specific ARGX-119 efficacy

Our *in vivo* experiments showed that ARGX-119 can reduce MuSK MG symptoms in a mouse model induced by one of the tested polyclonal patient IgG4, but not in experiments with three other patient IgG4s. To explain these patient-specific differences, we considered three possible mechanisms by which ARGX-119 efficacy could be limited in some patients. First, some patients may have (high affinity) MuSK Fz-domain binding antibodies which may compete for ARGX-119 binding. Fz-domain binding antibodies have been described in approximately 30% of MuSK MG patients.^17, 18^ Second, by binding to other MuSK domains, patient antibodies may alter MuSK conformation, preventing ARGX-119 from forcing dimerization. Lastly, bivalent agonistic patient-derived MuSK antibodies have been described and may lead to myasthenic symptoms.^21, 27, 35^ Therefore, we hypothesized that some patients may have agonistic MuSK antibodies which may activate the MuSK kinase in a suboptimal manner, preventing the therapeutic agonistic effect of ARGX-119.

To further investigate these hypotheses, we performed epitope mapping in ELISA and surface plasmon resonance (SPR) experiments to determine whether physical obstruction by patient 2, 3 and 4 antibodies played a role in the limited efficacy of ARGX-119 in our disease models. Epitope mapping revealed the dominant epitope to be the Ig-like domain 1 for all patients (Figure 3A). Patient 1 and 2 had additional antibodies against the Ig-like domain 2 but, given the therapeutic benefit in the model using patient 1 IgG4, these Ig-like domain 2 antibodies are likely not to affect ARGX-119 efficacy, arguing against the second hypothesis. Patient 2, in addition, had antibodies binding the Fz-domain. In SPR experiments, moderate competition with ARGX-119 (using human, but not mouse, MuSK) could be observed, suggesting that the lack of efficacy of ARGX-119 with this patient 2 material in our mouse experiments is unlikely due to competition (Figure 3B, C and Supplementary Figure 5). Fz-domain binding was not observed for the other patient materials. In conjunction with this, no competition with ARGX-119 was observed with IgG4 from patients 1, 3 and 4. Thus, lack of ARGX-119 therapeutic efficacy with three of the patient materials could not be explained by direct competition.

**Figure 3:**
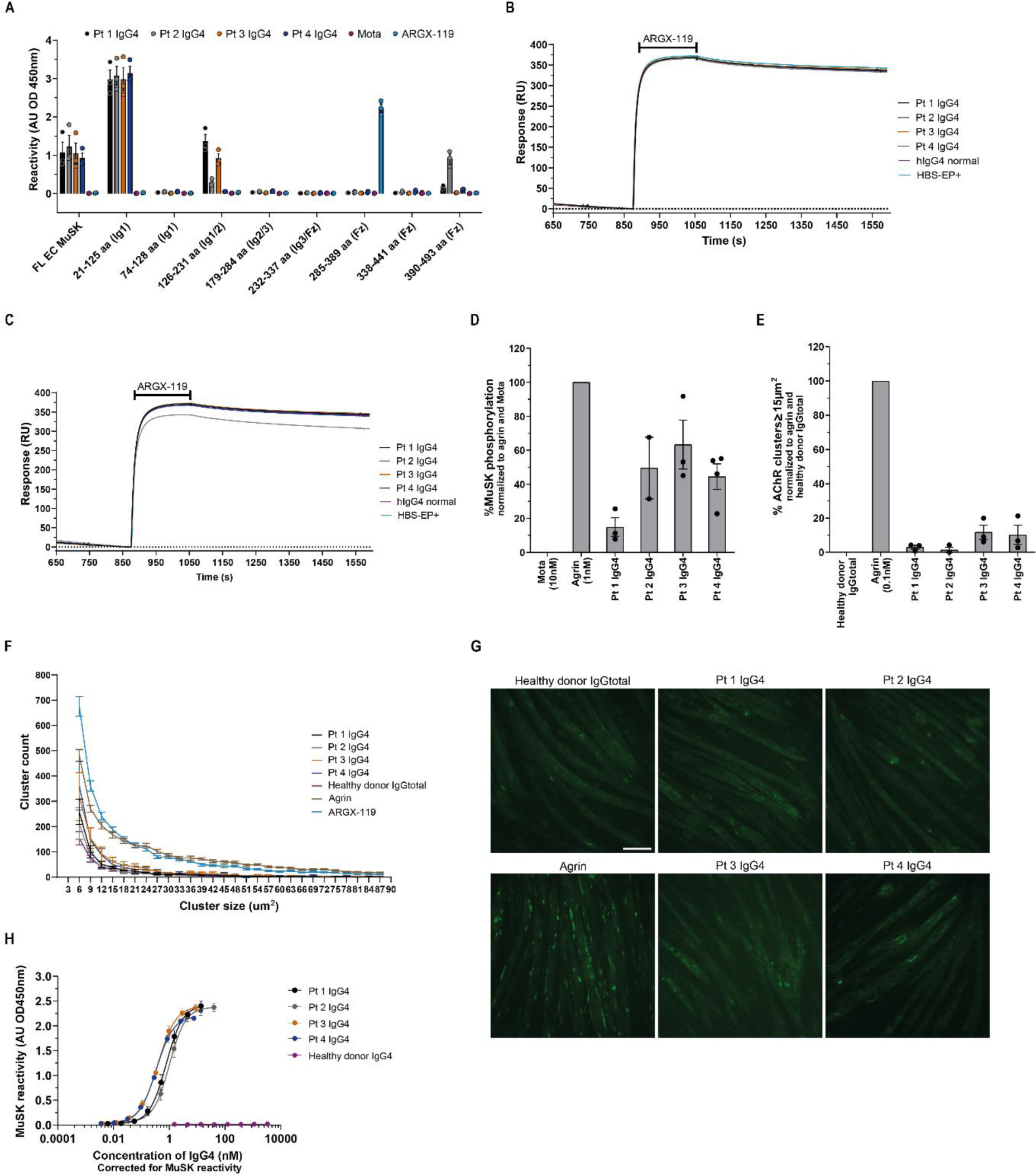
*In vitro* assessment of patient-specific ARGX-119 efficacy. A) Epitope mapping of pt IgG4s binding to different parts of human MuSK. B) Sensorgrams of the binding of patient IgG4 to mouse Fz-domain of MuSK in competition with ARGX-119. C) ) Sensorgrams of the binding of patient IgG4 to human Fz-domain of MuSK in competition with ARGX-119. D) Maximum endogenous agonistic potential of patient material on MuSK phosphorylation in C2C12 myotubes. Concentration of IgG4 per patient material: Pt 1 293.41 nM, Pt 2 146.75 nM, Pt 3 292.50 nM, Pt 4 293.48 nM. E) Maximum endogenous agonistic potential of patient material on AChR clustering in C2C12 myotubes. Concentration of IgG4 per patient material: Pt 1 730.72 nM, Pt 5.42 nM, Pt 3 34.91 nM and Pt 4 62.94 nM. F) AChR cluster size distribution of each conditions in D. G) Representative images of each condition in D (visualized by BTX488 staining). H) MuSK biobridging assay detecting bivalent MuSK binding antibodies. Data represent mean ± SEM. One-way ANOVA with Tukey multiple comparisons test (C and D). Scale bar: 100 µm.

Differentiated C2C12 myotube cultures contain all the necessary muscle-specific machinery of the postsynaptic neuromuscular junction, and by adding neural agrin and patient antibodies to the culture, they can be used to model MuSK MG *in vitro*.^21, 36, 37^ Exposure of these myotube cultures to patient antibodies, in the absence of the natural MuSK activator agrin allowed us to assess whether patient material has agonistic potential and may prevent or impair ARGX-119 induced dimerization in line with the third hypothesis described above. All patient IgG4s had some capacity to activate the MuSK kinase (Figure 3D), which also translated in increased AChR cluster numbers in two out of four patients (Figure 3E, F and G). The isotype control antibody Mota gave no MuSK kinase activation, nor AChR clustering. IgG4 from patient 1, the *in vivo* pathogenic material on which ARGX-119 showed therapeutic efficacy in mice, had the lowest level of MuSK activation (phosphorylation), i.e. ∼15% of the agrin level. Patients 2, 3 and 4 gave MuSK activation up to ∼45-63% of the agrin condition. Matching our previous observations, induction of MuSK phosphorylation did not translate linearly to AChR clustering, with AChR cluster levels close to the control healthy donor IgG condition for patients 1 and 2 and reaching ∼15% of the agrin condition in patients 3 and 4.^37^ AChR cluster size in myotube cultures is a measure of AChR cluster maturity. To investigate whether patient antibodies differentially affect AChR cluster size we performed cluster size distribution analysis which showed that IgG4 from patients overall induced lower numbers of AChR clusters of all sizes compared to agrin and ARGX-119 treatments (Figure 3F). IgG4 from patient 3 and 4 however, induced more smaller clusters than IgG4 from patient 1 and 2 and neared the small cluster size numbers of the agrin condition. These results suggest that some patients may carry MuSK activating antibodies which could prevent proper MuSK activation through ARGX-119.

In most MuSK MG patients, autoantibodies of the IgG4 type make up >90% of all MuSK reactive antibodies.^38^ However, some patients do have (bivalent) antibodies against MuSK of different subclasses which may stimulate MuSK activation.^27, 35, 38, 39^ To exclude that contamination of our purified IgG4 fractions by bivalent and potentially agonistic MuSK antibodies of other subclasses interfered in these assays, we determined the presence of other IgG subclasses in our purified IgG4 material (Supplementary Table 2). These data show that levels of combined IgG1, IgG2 or IgG3 in the purified IgG4 fractions was ∼6%, ∼6%, ∼9%, ∼10% respectively for patients 1-4. This suggests that, given the already low levels of MuSK autoantibodies in IgG1-3 fractions of MuSK MG patients^38^, an agonistic effect due to MuSK IgG1-3 antibodies is unlikely.

To further investigate whether some IgG4 fractions of MuSK MG patients unexpectedly have bivalent MuSK antibodies we developed a biobridging assay (Figure 3H and Supplementary Figure 6). In this assay, bivalent MuSK antibodies were detected by cross-linking a plate-bound MuSK and a soluble tagged MuSK protein. All patient IgG4 samples had some bivalent MuSK binding antibodies (Figure 3H) with patient 3 and 4 material having the highest concentration and the two others having equal levels of bivalent MuSK antibodies (EC50 patient 1-4: 0.77 - 1.01 – 0.39 – 0.34). This presence of bivalent MuSK antibodies in the IgG4 fraction is in line with the agonist activity of the purified IgG4 samples observed in Figure 3, but does not appear to explain the differential ARGX-119 efficacy *in vivo* by itself.

In conclusion, lack of efficacy of ARGX-119 observed with three of our MuSK MG patient IgG4s in the MuSK MG passive transfer mouse models could not be explained by direct competition for binding with ARGX-119, but rather seems related to a fraction of (IgG4) MuSK (bivalent) antibodies which are capable of inducing significant amounts of MuSK phosphorylation present in some patient materials which warrants further investigations.

### *In vitro* therapeutic rescue may be predictive for *in vivo* therapeutic efficacy of ARGX-119

A major challenge in clinical practice is to identify which patients will benefit from a certain type of treatment. Predictive screening assays could greatly reduce risk of exposure to ineffective treatments with associated side-effects for patients, and have the potential to lower health care costs for society.^40^ We therefore set out to develop a method to identify MuSK MG patients who are likely to benefit from therapeutic treatment with ARGX-119. We designed a C2C12 myotube assay to assess ARGX-119-based therapeutic effects on AChR clustering, in the presence of patient antibodies to model patient-specific disease (Figure 4A).

**Figure 4:**
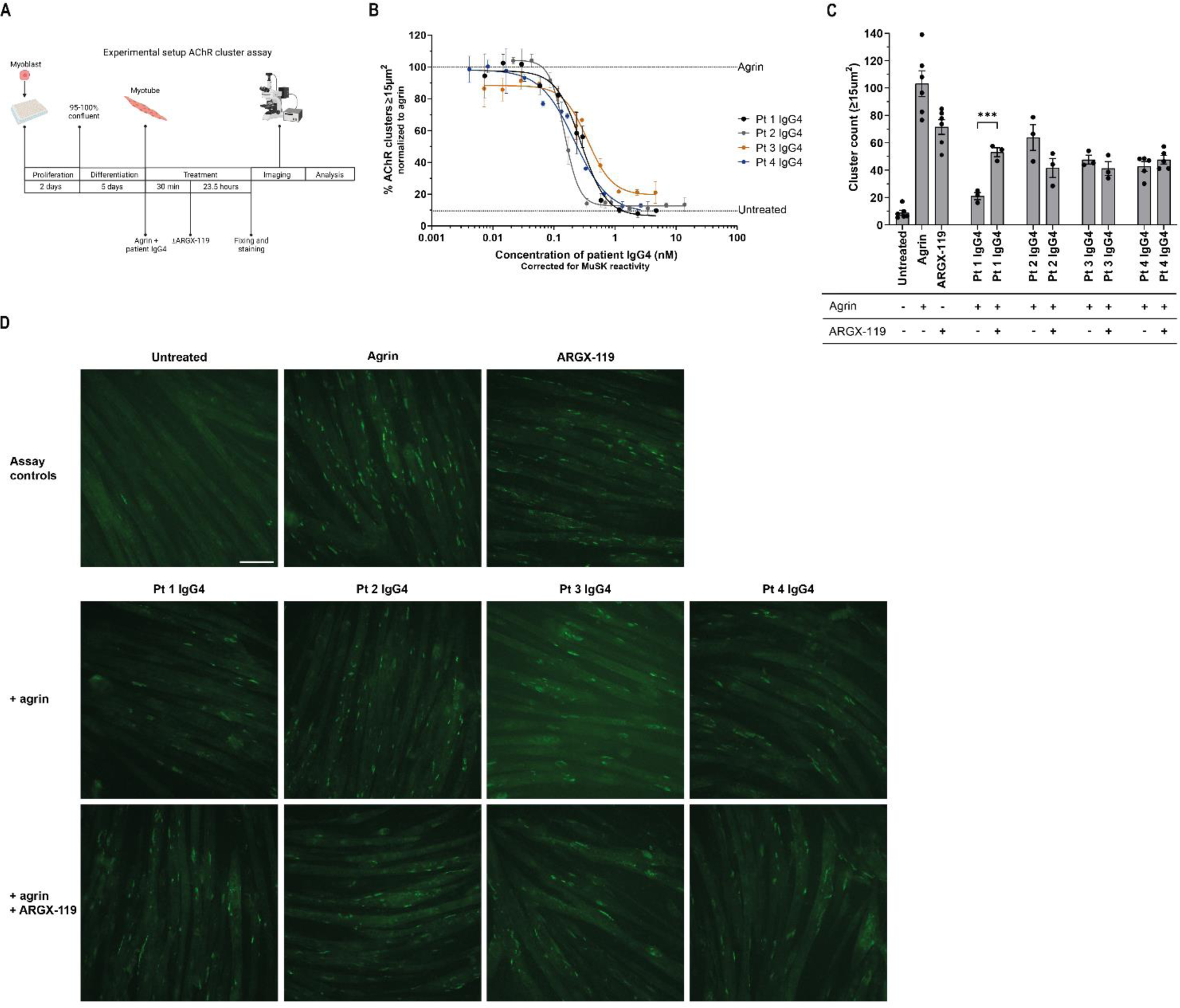
An *in vitro* therapeutic assay may be predictive for *in vivo* ARGX-119 efficacy in MuSK MG. A) Experimental setup of the AChR clustering assay. B) Inhibition potential of patient IgG4 on AChR clustering in C2C12 myotubes. C) Quantification and D) representative images of the therapeutic effect by ARGX-119 (0.1563 nM) on AChR clustering in C2C12 myotubes treated with patient IgG4. Data represent mean ± SEM. Paired t-test was used in C. *P < 0.05, **P < 0.01, ***P < 0.001. Scale bar: 100 µm.

MuSK autoantibody content and potency varied between patients^21, 41^ and largely matched the potency observed in the animal experiments (Figure 4, Supplementary Figure 1). To develop a personalized *in vitro* model that provides a therapeutic window for testing ARGX-119, we first determined the IC60 per patient material in these assays in the presence of agrin (Figure 4B). Next, we assessed the improvement of AChR clustering in the presence of ARGX-119. Clear improvement was observed with patient 1 IgG4, but not with the three other IgG4s (Figure 4C and D). Since ARGX-119 has a bell-shaped dose-effect relationship (Figure 2), we also tested several other doses to ensure we were not overlooking any ARGX-119 induced therapeutic effect (Supplementary Figure 7). These experiments confirmed that different doses of ARGX-119 did not result in *in vitro* improvement of AChR clustering when using IgG4s from the three *in vivo* non-responsive patients’ models confirming our *in vivo* therapeutic tests (Supplementary Figure 4). In conclusion, the *in vitro* therapeutic test results perfectly correlate with the therapeutic effects observed in the *in vivo* mouse experiments, suggesting that this method may be used to identify ARGX-119 sensitive MuSK MG patients. Further validation of the assay with clinical responses will be required for future clinical application.

## Discussion

MuSK MG is a potentially life-threatening disease that often requires chronic immunosuppression and responds poorly to symptomatic treatment with acetylcholine esterase inhibition (e.g. pyridostigmine).^42^ Rituximab, an anti-CD20 B cell depleting therapy, is often successfully used in MuSK MG patients. However, it comes with additional risks of infection and some patients become drug resistant.^43^ It is estimated that ∼10 % of patients do not respond sufficiently to the available therapies.^42^ Here, we present an alternative therapeutic strategy for MuSK MG, by targeting the trophic MuSK pathway directly with a MuSK agonist antibody, thus counteracting the inhibitory effect of the patients’ pathogenic MuSK autoantibodies. This might be a useful complementary therapeutical effect that cannot be obtained by any means of immunosuppressive treatment.

In the current study, the MuSK agonist antibody ARGX-119 ameliorated myasthenic disease in mice caused by injection of IgG4 antibodies from one of four MuSK MG patients. In addition, in a collaborative study, a therapeutic effect of ARGX-119 was observed in a mouse model of MuSK MG, caused by patient-derived functionally monovalent monoclonal MuSK antibodies (Personal communication Steve Burden, Neurology Department, Massachusetts General Hospital/Harvard, Boston, MA, USA). These studies showed that ARGX-119 improved the organization and function of neuromuscular synapses and rescued mice from motor performance deficits and ultimately lethality. Our study presented here takes a next step towards clinical application by testing ARGX-119 in a passive transfer mouse model using polyclonal antibodies from symptomatic MuSK MG patients. This IgG4 is complex and diverse in terms of epitopes they recognize, and agonistic or antagonistic effects on MuSK. The data in this study reveals heterogeneity in the MuSK MG population regarding the therapeutic potential of MuSK agonist ARGX-119 and suggests that the effect of ARGX-119 may depend on the nature of the individual MuSK autoimmune response.

The preclinical model described here made use of polyclonal patient-derived IgG4 to induce MG in mice. Antibodies against MuSK of other subclasses may also have pathogenic potential *in vitro*.^21, 35^ However, pathogenicity was not observed when performing passive transfer with IgG1-3 fractions from MuSK MG patients.^14^ This is probably related to the low amount of MuSK autoantibodies of the IgG1-3 type present in patients serum.^38^ Passive transfer with total IgG therefore poses a technical challenge as sufficiently high doses of MuSK antibodies cannot always be administered to mice to induce a phenotype. In light of this, the predominance of MuSK autoantibodies often being well over 90% of all MuSK antibodies in patients, and the successful use of this model in previous preclinical test of e.g. efgartigimod (an FcRn inhibitor), which is currently on the market to treat MG patients,^28, 44^ the use of IgG4 to model MuSK MG in mice was considered the best option for these preclinical tests.

Mechanistic studies looking into the inter-patient differences in ARGX-119 efficacy showed that there was no correlation between pathogenic potency of patient IgG4 and ARGX-119 efficacy. Furthermore, one patient had Fz-domain binding antibodies which did not compete for ARGX-119 binding to mouse MuSK in SPR studies. Although in our experiments competition for ARGX-119 binding was not observed and could not explain lack of ARGX-119 efficacy in three patients’ models, this mode of inhibition of this therapeutic strategy may still be relevant for ∼30% of MuSK MG patients who carry relatively large amounts of Fz-domain autoantibodies.^17, 18^ In addition, the three patient IgG4s for which no therapeutic effect of ARGX-119 was observed in the mice, had a relatively large agonistic antibody fraction which induced MuSK phosphorylation and for some translated to AChR clustering independent of the natural ligand agrin. We hypothesize that these high affinity MuSK agonistic autoantibodies in the patient serum force MuSK dimerization in a suboptimal manner as previously described, preventing the proper activation of MuSK by ARGX-119 (Figure 5).^37^ This may suggest that MuSK MG patients who have (large amounts of) agonist autoantibodies, may not benefit from ARGX-119 treatment. An alternative explanation could be that some of these autoantibodies activate the MuSK kinase while blocking other essential roles of MuSK like clustering of Lrp4 for maintaining presynaptic differentiation, which can likely not be rescued by a MuSK agonist. This seems however unlikely since the four tested patient materials had a similar epitope pattern and dominance of the Ig-like 1 domain response while showing differential efficacy of ARGX-119.

**Figure 5:**
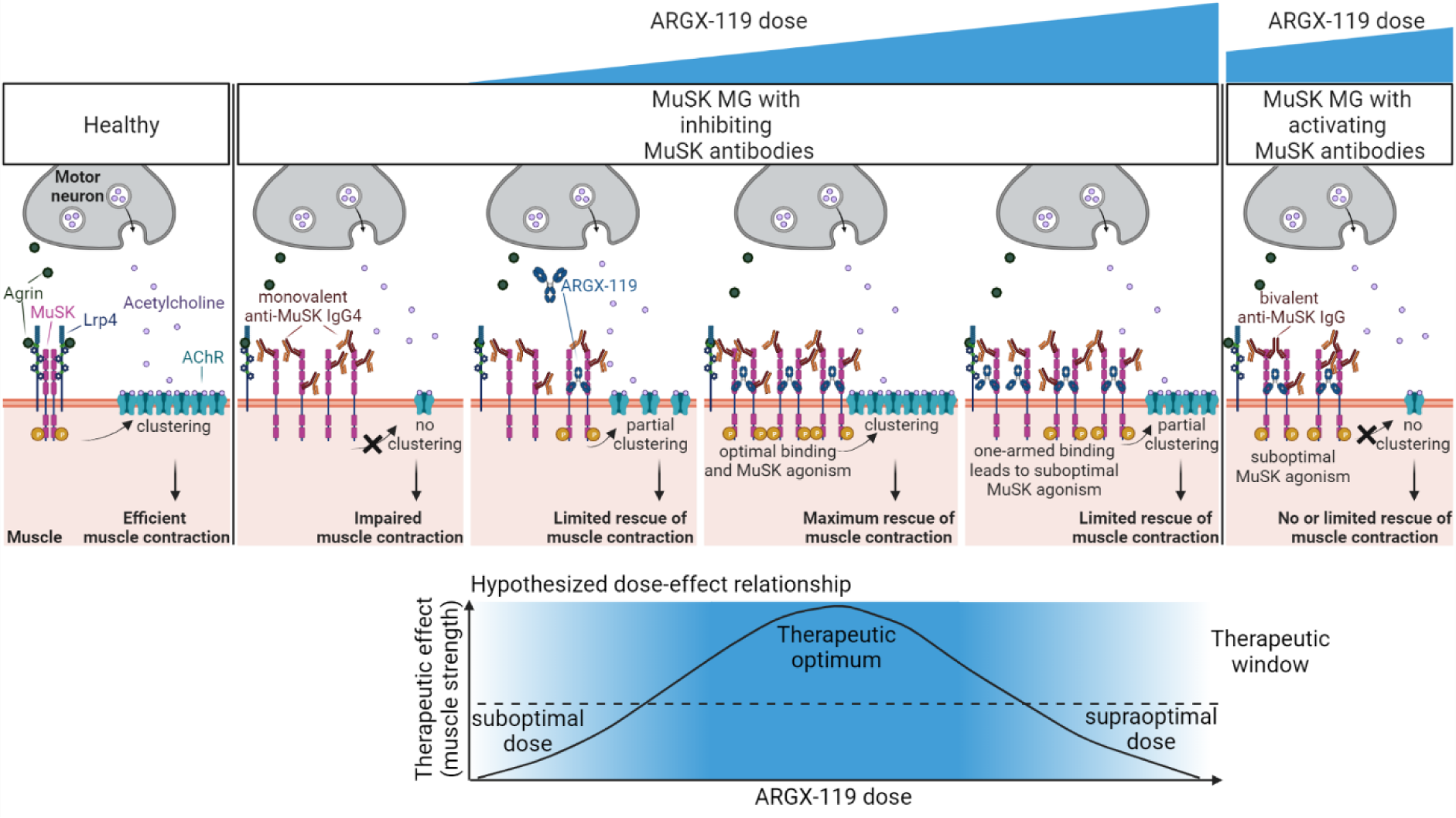
Schematic visualization of the hypothesized mechanisms underlying the patient-specific therapeutic benefit of MuSK agonist antibody ARGX-119 in MuSK MG passive transfer models. Neuromuscular synapses in health are established and maintained by the agrin-Lrp4-MuSK pathway. In MuSK MG monovalent anti-MuSK IgG4s inhibit activation of MuSK by agrin-Lrp4. Suboptimal doses of agonistic MuSK antibodies such as ARGX-119 do not dimerize and thereby activate enough MuSK molecules to sufficiently drive AChR clustering and have a therapeutic effect. At the therapeutic optimum there is enough ARGX-119 to dimerize and activate MuSK to maximally drive AChR clustering which has the most therapeutic effect. At the supra-optimal dose, there is too much ARGX-119 for the amount of available MuSK molecules. This results in monovalent binding of ARGX-119 to MuSK and thereby not enough dimerization and activation of MuSK, and limited therapeutic benefit. When a MuSK MG patient has a significant amount activating MuSK autoantibodies, they may force MuSK in a suboptimal structural conformation, perhaps blocking other essential roles of MuSK and preventing therapeutic MuSK agonism by ARGX-119, independent of its dose.

These findings furthermore prompted us to seek a screening assay to identify those MuSK MG patients who might benefit from treatment with ARGX-119. We found that the ability of ARGX-119 to ameliorate disease *in vivo* correlates with the ability to counteract patient autoantibody-mediated inhibition of AChR clustering in cultured muscle cells. Due to limited amounts of these rare plasmapheresis materials (these *in vivo* studies requiring liters of material) we were however unable to extend the passive transfer experiments with more unrelated patients materials and confirm the hypothesis in this study. It will be important to investigate how predictive both the *in vitro* and *in vivo* studies are for a therapeutic effect of ARGX-119 in MuSK MG patients. The simple and rapid *in vitro* screening assay has the potential to lower the risk of exposure to ineffective treatments for these patients, as well as lower health care costs for society.

MuSK agonism can be achieved by a variety of bivalent recombinant antibodies targeting different epitopes on MuSK. Patient-derived bivalent MuSK antibodies binding the Ig-like 1 domain can induce MuSK phosphorylation and (partial) AChR clustering but did not improve muscle weakness in a MuSK MG passive transfer model using purified patient IgG4 (the same patient 1 IgG4 used in this study).^37^ The Ig-like 1 domain antibodies furthermore induced an unexpected and unexplained mouse urological syndrome which led to sudden death in a proportion of the used male mice, specifically of the C57BL/6 strain. Urological side effects have thus far not been reported with Fz-domain binding MuSK antibodies in several preclinical studies (Vanhauwaert et al. 2024. bioRxiv doi: 10.1101/2024.07.18.604166).^32^ The Ig-like 1 domain of MuSK is an important interaction region for several other neuromuscular synaptic proteins including Lrp4, Vangl2, agrin and biglycan.^11, 19, 45, 46^ Lrp4 furthermore anchors collagen Q and AChE to the post-synaptic membrane.^47^ The MuSK Fz-domain may interact with Lrp4 and wnt factors,^48, 49^ but is dispensable in mice as mice lacking the MuSK Fz-domain are viable and have normal neuromuscular synapses.^50^ We therefore hypothesize that MuSK agonism via antibodies binding the Ig-like 1 domain interferes with some of these other functions of MuSK, whereas Fz-domain binders are devoid of this and only stimulate MuSK signalling when this is perturbed by autoantibodies or otherwise. Importantly, MuSK activation and phosphorylation follows a particular pattern.^51^ Successful MuSK agonism, next to not interfering with endogenous MuSK function, should induce downstream signalling similar to the natural ligand agrin.^52^ It will be interesting to learn what characteristics of a MuSK antibody define a successful therapeutic agonist. As described for agonistic antibodies of dimerizing membrane ligands other than MuSK,^34^ we also see a bell-shaped curve in most of the metrics tested using ARGX-119 (Figure 5). Other studies testing ARGX-119 in different neuromuscular disease models however showed efficacy with doses exceeding the optimal dose in our experiments (Vanhauwaert et al. 2024. bioRxiv doi: 10.1101/2024.07.18.604166). This suggests that careful dose finding should be done in the early clinical phase and that doses might be different across indications. Indeed, synaptic MuSK levels have been described to vary between muscles from different neuromuscular disease models. For example, MuSK expression levels were greatly reduced in a mouse model for Duchenne muscular dystrophy, a monogenetic muscle wasting disease in which neuromuscular synapse abnormalities have been described.^53–55^ Similarly, levels are reduced in mice exposed to MuSK MG patient IgG^29^ or after nerve crush damage.^56^ On the other hand, MuSK levels may be normal in other neuromuscular diseases associated with neuromuscular synaptic deficits, such as SMA.^57^ MuSK agonist treatment may thus have a disease-specific optimal dose-range.

MuSK is a key regulator of neuromuscular synaptic stability. The trophic signalling pathway converging on MuSK has been assumed to have therapeutic potential for a variety of diseases characterized by or associated with neuromuscular synaptic defects,^58^ especially to counteract muscular atrophy. These diseases include ALS, SMA, sarcopenia and other forms of (congenital) MG. Although in most of these diseases a MuSK agonist will not resolve the underlying pathophysiology, improved neuromuscular synaptic stability and transmission may maintain muscle performance and innervation, and thereby improve muscle function or prevent progression in these diseases. MuSK agonism using different Fz-domain binding antibodies (X17 or Mab13) have so far improved symptoms of muscle weakness in animal models for genetic neuromuscular disorders such as SMA, ALS and Dok7 CMS.^31, 32, 59^ Our study now expands this list by showing therapeutic potential of a new Fz-domain targeting MuSK agonist in a model for an acquired autoimmune disease. It will be exciting to learn whether other neuromuscular diseases will benefit from ARGX-119.

To conclude, this study translates the therapeutic potential of ARGX-119 one step closer to MuSK MG patients, showing that 1. Efficacy has a bell-shaped dose-effect relationship in MuSK MG, 2. Efficacy may be limited to a subset of MuSK MG patients, and 3. These patients may be identified with an *in vitro* screening assay.

## Materials and methods

### Study approval

To develop the MuSK MG mouse models, antibodies were isolated from therapeutic plasmapheresis material from MuSK MG patients. Patients participating in this study were recruited at the MG outpatient clinic of Leiden University Medical Center (LUMC) and signed informed consent. No permission from a medical ethical committee was needed for the use of this “waste” material.

Animal experiments were carried out according to Dutch law and Leiden University guidelines, including approval by the local Animal Experiments Committee (Instantie voor Dierenwelzijn).

### Antibody isolation, production and selection

Frozen (-80°C) plasmapheresis materials from four unrelated patients, were selected for IgG4 purification, based on high MuSK reactive IgG4 titers and presence of large volumes required for the *in vivo* experiments. The different batches per patient were transported to Immunoprecise Antibodies Ltd (Utrecht, the Netherlands), pooled and purified for IgG4 on a CaptureSelect IgG4XK20/50 column (Pharmacia Biotech). The concentrated IgG4 in PBS buffer was assessed for purity via LabChip analysis using the Protein Clear HR kit (Revvity) and indicated a minimal purity of ∼95% for all the patient IgG4 pools, and shipped back to the LUMC for storage at −20°C until further use. Patient characteristics are detailed in Supplementary Table 1.

We previously produced and characterized a Llama-derived Fz-domain binding MuSK antibody (ARGX-119) which showed strong MuSK agonistic potential and rescued a Dok7 CMS phenotype in mice (Vanhauwaert et al. 2024. bioRxiv doi: 10.1101/2024.07.18.604166). The here described preclinical therapeutic tests made use of ARGX-119 (3B2g2m1-hIgG1-LALAdelK) and isotype control (Mota-hIgG1-LALAdelk) (Mota), which were produced by Lonza Biologics (Slough, UK) and Immunoprecise Antibodies Ltd (Utrecht, The Netherlands). Mota, the isotype control antibody, is an anti-Respiratory Syncytial Virus F glycoprotein (RSV-F) binding monoclonal antibody formatted in the same backbone as ARGX-119 and cannot bind to MuSK.

### Mouse passive transfer studies

To test the therapeutic efficacy of ARGX-119 in a passive transfer mouse model with patient polyclonal IgG4, female NOD/SCID mice aged approximately 2 months at the start of each experiment were used. After establishing baseline values for the *in vivo* clinical signs during a training period of approximately 7 days, MuSK MG was induced by daily intraperitoneal (i.p.) administration of purified polyclonal IgG4 from either of the four unrelated patients. Mice were randomized across treatment groups based on starting weight to create homogenous groups. During the experiment, mice were assessed for general wellbeing, body weight (Ohaus CS 200 scale), grip strength and hanging time on an inverted mesh. Forelimb and abdominal muscle strength was assessed using a grip strength meter (type 303500, Technical and Scientific Equipment GmbH). The average peak force value of a trial of 10 consecutive pulls, with a few seconds pause, was calculated. The inverted mesh hanging test was used to assess fatigability of limb and abdominal muscles as described previously.^60^ The test ended upon completing the maximum hanging time of 180 seconds. If a mouse fell before reaching the 180 seconds, the test was repeated up to three times. The best of three attempts was used in the data analysis. Body weight and grip strength data were normalized to day 0, just before the first IgG4 injection. If mice had body weight loss of ≥20% of the starting weight, their humane endpoint was considered reached, and mice were euthanized by CO2 inhalation.

Potency of patients IgG4 to induce MG varies significantly.^14^ We therefore tested in total four unique patients in this study. Before therapeutic tests were initiated, disease dose-finding experiments were carried out for each patient material to determine the minimal disease dose required for a clear myasthenic phenotype. For purified IgG4 from patient 1, the disease dose was previously published.^37^ To estimate the dosing range to be tested in the dose-finding experiment for the other patients, we compared MuSK reactivity levels in a MuSK ELISA^17^ and related that to our historic experience with disease doses from other patient material and their MuSK reactivity levels.^14, 37^

For therapeutic tests, the disease-causing dose per patient IgG4 was administered to the mice daily. Different doses of ARGX-119 (20, 10, 2.5, 0.25, 0.1 or 0.01 mg/kg) or isotype control antibody Mota (10, 2.5 or 0.25 mg/kg) were injected i.p. every 3 days and initiated one day prior to the first injection of polyclonal patient IgG4 and co-injected thereafter. Injections were administered until the humane endpoint or the end of the experiment was reached (Figure 1A). The experimenter was blinded to the treatment.

Blood samples for analyzing serum titers were taken through tail vein puncture one day prior to the start of injections and on the days of ARGX-119/isotype control antibody (and IgG4) injections (shortly before). All blood samples were spun down at 2000 rpm for five minutes, and serum was subsequently stored at -20°C until further use.

### Serum titer analysis of ARGX-119 and IgG4 patient antibodies

To determine pharmacokinetics of ARGX-119 in mouse serum, we quantified the concentrations using a qualified ELISA method. 1 µg/mL anti-idiotypic antibody (mouse IgG1, in-house production) in 1xPBS, was coated overnight at 4°C on a MaxiSorp plate (ThermoFisher Scientific). The next day the assay plate was blocked with 1% casein blocking buffer (Bio-Rad) for one hour at room temperature. A calibrator curve using ARGX-119 (ImmunoPrecise Antibodies Ltd, Utrecht, The Netherlands) was prepared in C57BL/6 gender-pooled mouse serum (BioIVT, cat MSE01SRM) in assay buffer (0.1% Casein in 1xPBS). Study samples were also diluted in assay buffer and then further diluted in sample buffer (0.1% Casein in 1xPBS and 1% serum). The calibrator and sample dilutions were added in duplicate to the assay plate and incubated for one hour at room temperature. ARGX-119 was detected using a biotinylated anti-LALA hFab (in-house production) in assay buffer for one hour at room temperature followed by adding HRP-conjugated streptavidin in assay buffer for 30 minutes at room temperature. The color development reaction, using s(HS)TMB substrate, was terminated by adding sulfuric acid and optical density was measured at 450 nm (reference 620 nm) on an Infinite M Nano plate reader (Tecan). Results were processed using Graphpad Prism 9 software.

To confirm mice were equally exposed to patient antibodies we quantified patient IgG4 levels in the mouse serum by means of ELISA. 1 µg/mL anti-human IgG4 monoclonal ab (clone HP6023, ThermoFisher Scientific) in 1xPBS was coated overnight at 4°C on a MaxiSorp plate (ThermoFisher Scientific). The next day the assay plate was blocked with 1% casein blocking buffer (Bio-Rad) for one hour at room temperature. A calibrator curve using the affinity purified patient polyclonal IgG4 (ImmunoPrecise Antibodies Ltd, Utrecht, the Netherlands) was prepared in serum and assay buffer (0.1% Casein in 1xPBS). Per study the injected patient IgG4 was used as calibrator. Study samples were also diluted in assay buffer and then further diluted in sample buffer (0.1% Casein in 1xPBS and 1% serum). The calibrator and sample dilutions were added in duplicate to the assay plate and incubated for two hours at room temperature. Patient IgG4 was detected using 25 ng/mL of an HRP-conjugated goat F(ab’)2 anti-human IgG-Fc polyclonal antibody (Abcam) in assay buffer for one hour at room temperature followed by adding TMB substrate. Color development reaction was terminated by adding sulfuric acid and optical density was measured at 450 nm (reference 620 nm). The plate was analyzed on an Infinite M Nano plate reader (Tecan). Results were processed using Graphpad Prism 9 software.

### Epitope mapping and binding assays

For epitope mapping of patient IgG4 antibodies, an ELISA was performed with MuSK protein fragments as described previously.^17^

### SPR experiments

To assess for competition of patient IgG4 antibodies against ARGX-119, surface plasmon resonance (SPR) experiments were done with the Fz-domain of human or mouse MuSK.

A CM5 chip (Cytiva, cat 29149604) was immobilized with StrepmAb-IMMO (IBA-Lifesciences, cat 2-1517-001) using amine coupling on a Biacore 3000 SPR device (GE, cat BR110045). In-house produced human or mouse Fz-domain of MuSK, was captured, followed by MG patient purified IgG4s and subsequently ARGX-119. Binding response (in RU) was measured just before termination of the injection of patient IgG4s or ARGX-119. For all steps HBS-EP+ pH7.4 was used as running buffer.

### C2C12 myotube cultures

C2C12 myoblasts were obtained from CLS Cell Line Service. They were maintained in Dulbecco’s Modified Eagle Medium containing GlutaMAX (10569-010, Gibco) and supplemented with 10% fetal bovine serum (S-FBS-CO-015, Serana) and Penicillin Streptomycin (15140-122, Gibco) at 37C°. For MuSK phosphorylation studies, cells were plated in 6-well plates and differentiated in medium supplemented with 1% horse serum (H1138, Sigma) and Penicillin-Streptomycin (P4333, Sigma). For AChR clustering studies, the myoblasts were cultured in 96-well plates (Greiner µclear, 655090, 0.32 cm^2^/well) in proliferation medium until 90-95% confluency. Next, the proliferation medium was replaced with low serum differentiation medium containing 2% horse serum (HI, 26020-088, Gibco), which was changed every other day until mature myotubes had formed (∼5 days).

### MuSK phosphorylation

To investigate the agonistic potential of the different patient IgG4s, we performed a 2x serial dilution of the patient IgG4 in the absence of the natural ligand (agrin). Starting concentrations per patient IgG4 were 2348 nM. After 30 minutes incubation, cells were lysed and extensively scraped. Phosphorylated and total MuSK was detected in the lysates by means of a binding ELISA. MuSK-Ig1 ab clone 13-3B5-hIgG4 (Immunoprecise Antibodies) was immobilized onto the assay plate and the lysates were loaded in quadruplicate, which allowed for detection of both total MuSK, via incubation with MuSK-Ig1 ab clone 3F6c (Immunoprecise Antibodies, in-house biotinylated), and tyrosine phosphorylated MuSK, via incubation with a mix of bioinylated phosphotyrosine abs clone 4G10 (16-103, Merck Millipore Corp) and clone PY20 (309304, Biolegend) in the same in duplicate. Bound antibodies were detected via poly-HRP conjugated streptavidin protein (PIER21140, Pierce) followed by a coloring reaction using TMB substrate. MuSK phosphorylation signals were normalized to the total MuSK signal (P-MuSK/MuSK). The quantified MuSK phosphorylation levels were normalized to the phosphorylation induced by agrin (agrin = 100%) and by the isotype control (Mota = 0%). Finally, the concentrations were corrected for MuSK reactivity by the formula: (conc * %MuSK reactivity)/100. The experiment was repeated at least three times. Quantification of MuSK phosphorylation levels was performed as described previously (Vanhauwaert et al. 2024. bioRxiv doi: 10.1101/2024.07.18.604166).

### AChR clustering

For the dose-finding experiments of patient IgG4s, C2C12 myotubes were treated with rat agrin (0.11 nM, 550-AG, R&D systems) and a 11-point dilution series of each patient material ranging from 1.80 to 1150.33 nM and incubated for 24 hours at 37°C. Finally, the concentrations were corrected for MuSK reactivity by the formula: (conc * %MuSK reactivity)/100. For the therapeutic experiments with ARGX-119, myotubes were first treated with the 60% inhibitory concentration (IC60), based on the dose-finding of patient IgG4, for 30 minutes at 37°C. Then 11-point dilution series of ARGX-119 ranging from 0.0024 to 100 nM was added to the agrin and patient IgG4 and incubated for 23.5 more hours, and as controls untreated, agrin (0.11 nM) and ARGX-119 (2.5 nM) were used. For the experiments testing the agonistic potential of the different patient IgG4s, myotubes were treated with healthy donor IgGtotal (1568.63 nM), agrin (0.11 nM) or different ICs of patient material [IC5, IC15, IC30, IC60, IC90 and IC100], which incubated for 24 hours at 37°C. After the incubation described above, cells were fixed in 4% w/v paraformaldehyde for 10 minutes at room temperature, and then incubated with Alexa Fluor 488-conjugated α-bungarotoxin (B13422, Thermo Fisher) to stain AChRs and Hoechst 33342 (H1399, Thermo Fisher) to stain nuclei, for 30 minutes at room temperature. For imaging of AChR clusters, a Leica AF6000 fluorescence microscope was used at 20x magnification. Five visual fields per well and four wells per condition were selected, based on evenly spread and clearly visible mature myotubes in the brightfield channel. The analysis of AChR-positive clusters count was done in ImageJ (1.52n), and the number of mature clusters (≥15 µm^2^) per condition was quantified. All experiments were repeated at least 3 times.

### Bio-bridging ELISA

To assess whether the patient IgG4s possess bivalent MuSK antibodies, a bio-bridging ELISA was performed. Thereto, recombinant extracellular domain of human MuSK (ECD hMuSK) was coated overnight at a concentration of 1 µg/mL on a 96-well high binding half area plate (Greiner). The following day, the plate was washed and subsequently blocked with 1% casein blocking buffer (Bio-Rad). After blocking, the plate was washed four times with PBS-T (PBS with 0.05% Tween20) and subsequently dilutions of patient IgG4’s and controls, prepared in assay buffer (0.1% Casein in 1xPBS) were incubated. Unbound antibodies were removed with four washes using PBS-T, after which 1 µg/mL biotinylated ECD hMuSK was added. After one hour the plate was washed and bound MuSK was detected using 50 ng/mL streptavidin poly-HRP (Pierce). The plate was again washed four times with PBS-T and incubated with s(HS)TMB (SDT reagents). The reaction was stopped after approximately eight minutes with sulfuric acid (Chem-Lab). Absorbance (OD 450nm minus reference OD620nm) was measured on an Infinite M Nano plate reader (Tecan) and results were processed using Graphpad Prism 9 software. All incubations were done for one hour at room temperature unless otherwise indicated.

### Statistical analysis

Statistical analyses were performed using Graphpad Prism 8. The *in vivo* mouse model data was analyzed with a one-way ANOVA per timepoint with Dunnett’s multiple testing correction to compare clinical parameters between the ARGX-119 and Mota groups, except for percentage survival rates for which a Log-rank Mantel-Cox statistical test was used. Statistical analyses of in *vivo* data were only performed on average data points when both groups contained three or more living mice. Missing data (due to mouse sacrifice) in these experiments were imputed using the last value of the mouse until the end of the experiment (last observation carried forward approach). Statistical analysis of *in vitro* endogenous agonistic potential experiments used one-way ANOVA with Tukey multiple comparisons test to compare the effect of the different patient material. Analysis of the *in vitro* therapeutic experiments used paired t-tests. A p-value <0.05 was considered significant in the statistical analyses.

For determining the bell-shaped effect curve, maximum efficacy of different doses of ARGX-119 to decrease body weight loss and increase grip strength and hanging time was determined per individual animals by identifying the minima of individual time-vs.-effect curves using Certara Phoenix, non-compartmental analysis, model type “Drug Effect (220)”.

## Acknowledgements

This project was financed by argenx and Prinses Beatrix Spierfonds grant W.OR19-13. The LUMC forms part of the European Reference Network for Rare Neuromuscular Diseases (ERN EURO-NMD) and the Netherlands Neuromuscular Disorders Center (NL-NMD). MGH receives support from the LUMC Gisela Thier Fellowship 2021 and a 2019 ZONMW VENI.

We would like to thank Steve Burden and Julien Oury who provided valuable feedback throughout our experiments and Roy Augustinus and Manon van den Bout for preparatory work. We would like to thank the patients for donating material for this study.

## Author contributions

J.L.L collected, analyzed and interpreted the data and wrote the paper; S.M.J. collected, analyzed and interpreted the data and wrote the paper; J.J.P. conceived and designed the analysis, collected, analyzed and interpreted the data and critically revised the paper for important intellectual content; B.v.K. conceived and designed *in vitro* analysis, collected, analyzed and interpreted the data and critically revised the paper for important intellectual content; C.K. analyzed and interpreted the data and critically revised the paper for important intellectual content; R.C. collected, analyzed and interpreted the data and critically revised the paper for important intellectual content; C.S. collected, analyzed and interpreted the data, contributed analysis tools and critically revised the paper for important intellectual content; K.M. collected, analyzed and interpreted the data and critically revised the paper for important intellectual content; L.d.C. collected, analyzed and interpreted the data and critically revised the paper for important intellectual content; M.R.T. contributed to the acquisition and analysis of data and critically revised the paper for important intellectual content; P.U. contributed to the conception and design of the work, analyzed and interpreted the data and critically revised the paper for important intellectual content; K.S. contributed to the conception and design of the work, analyzed and interpreted the data and critically revised the paper for important intellectual content; S.M.v.d.M. contributed to the conception and design of the work, analyzed and interpreted the data and critically revised the paper for important intellectual content; D.L.E.V. contributed to the acquisition and analysis of data and critically revised the paper for important intellectual content; R.V. contributed to the conception and design of the work, analyzed and interpreted the data and critically revised the paper for important intellectual content; J.J.V. contributed to the conception and design of the work, analyzed and interpreted the data and critically revised the paper for important intellectual content; M.G.H. contributed to the conception and design of the work, contributed to the acquisition and analysis of data, analyzed and interpreted the data and wrote the paper.

