## Supplementary Appendix for "Patient-specific therapeutic benefit of MuSK agonist antibody ARGX-119 in MuSK myasthenia gravis passive transfer models"

### Supplementary data

#### Supplementary Table 1: Patient characteristics

| Patient IgG4 | Plasmapheresis material composition | Sex | Age at MuSK MG onset | Age at time of plasmapheresis | MGFA grade at time of plasmapheresis | Treatment | MuSK reactivity (nM) |
|---|---|---|---|---|---|---|---|
| 1 | Pooled from two plasmaphereses | M | 30 | 53 & 57 | 3B | Pyridostigmine, Prednisone, Mycophenolate | 182 |
| 2 | Pooled from two plasmaphereses | M | 19 | 22 | 2B | Pyridostigmine, Prednisone, Azathioprine, IvIgG (unsuccesful) | 607 |
| 3 | One plasmapheresis session | M | 67 | 69 | 3B | None | 142 |
| 4 | One plasmapheresis session | M | 53 | 53 | 2 | Prednison Azathioprine | 99 |

**Supplementary Table 2: Quantification of IgG subclasses in purified IgG4 batches quantified as a percentage of the sum of the subclasses.**

| | **IgG1 (% of sum of subclasses)** | **IgG2 (% of sum of subclasses)** | **IgG3 (% of sum of subclasses)** | **IgG4 (% of sum of subclasses)** |
|---|---|---|---|---|
| **Pt 1** | 3 | 3 | 0,1 | 94 |
| **Pt 2** | 3 | 3 | 0,1 | 94 |
| **Pt 3** | 7 | 2 | 0,1 | 91 |
| **Pt 4** | 4 | 6 | 0,1 | 90 |

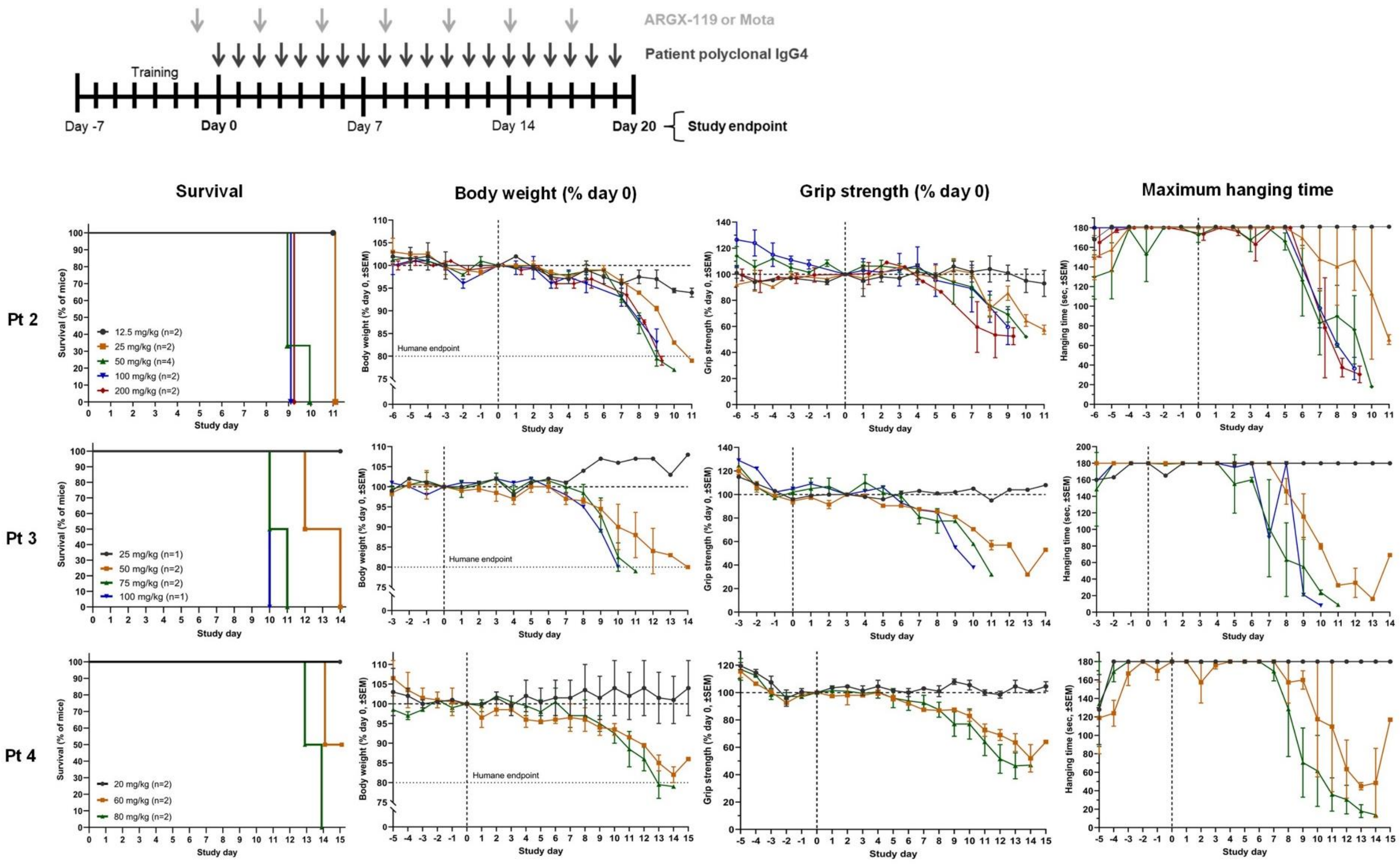


**Supplementary Figure 1: MG phenotype dose findings of purified patient IgG4.** *From left to right for patients 2-4*: percentage mouse survival over time, with reductions in survival representing sacrifice of mice due to humane endpoint reached; percentage body weight normalized to day 0; percentage grip strength normalized to day

0; and hanging time (maximum 180 seconds, best of 3 attempts) in mice treated with different doses of each purified patient IgG4. The doses in orange were identified as the minimal dose for a maximal MG phenotype per patient material. Data points of percentage grip strength and hanging time represent grouped values of the mean ±SEM of mice.

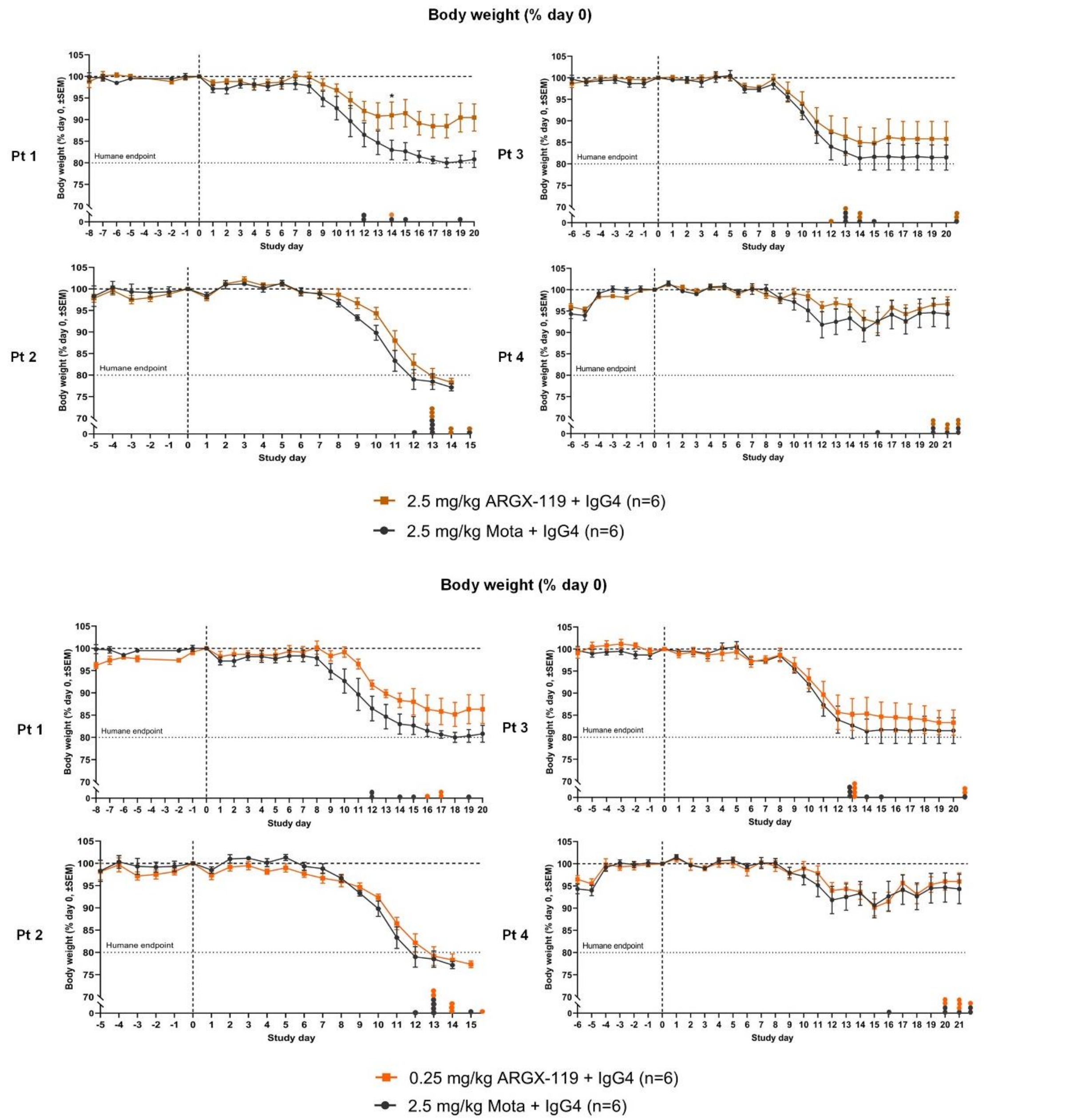


**Supplementary Figure 2: ARGX-119 improves body weight in the MuSK MG model induced with polyclonal patient IgG4 in a patient-specific manner.** Percentage body weight normalized to day 0 in mice treated with 2.5 or 0.25 mg/kg ARGX-119 or 2.5 mg/kg isotype control Mota. Data points of percentage body weight represent grouped values of the mean ±SEM of mice. Data points on the x-axis represent mice that were sacrificed as they reached humane endpoint or end of study. Dotted horizontal line represents the humane endpoint (80% body weight of day 0). Statistical analyses were only performed on average data points when the groups contained more than three mice. *$P<0.05$ for ARGX-119 vs Mota treated mice.

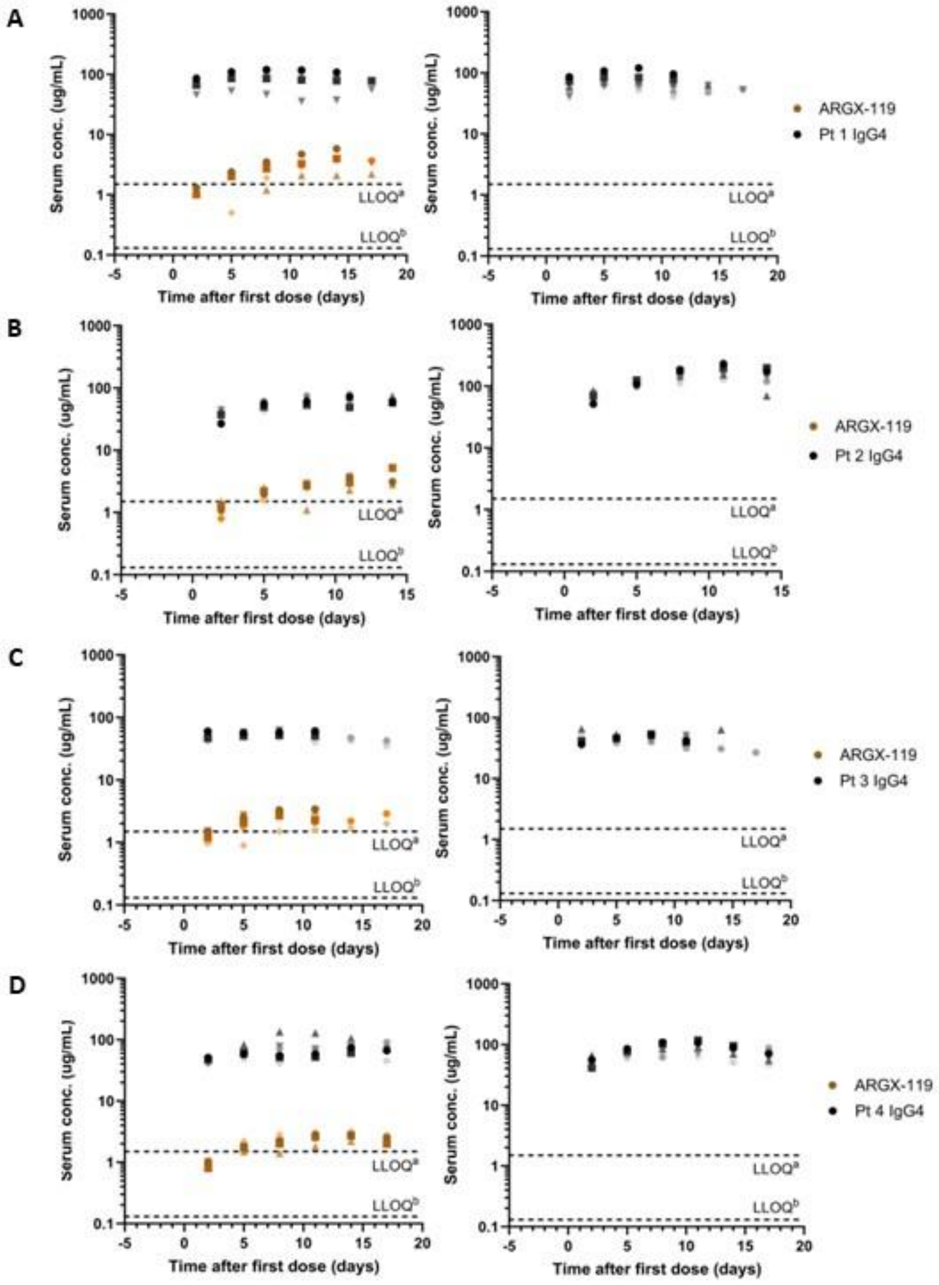


**Supplementary Figure 3: Exposure to ARGX-119 and polyclonal patient IgG4 was overall consistent between mice within the different experiments.** Serum concentrations (ug/mL) in individual mice over time until humane endpoint or end of study before and after administration of 0.25 mg/kg ARGX-119 or 2.5 mg/kg isotype control Mota (from left to right) and A) 40 mg/kg/day patient 1 IgG4, B) 25 mg/kg/day patient 2 IgG4, C) 50 mg/kg/day patient 3 IgG4 or D) 60 mg/kg/day patient 4 IgG4. Brown-to-orange colored data points represent ARGX-119 exposure in individual mice (each mouse represented by one color) and black-to-grey colored data points represent patient IgG4 exposure in the same mice (each mouse represented by one color). LLOQ[a], lower limit of quantification of patient IgG4 PK; LLOQ[b] of ARGX-119 PK.

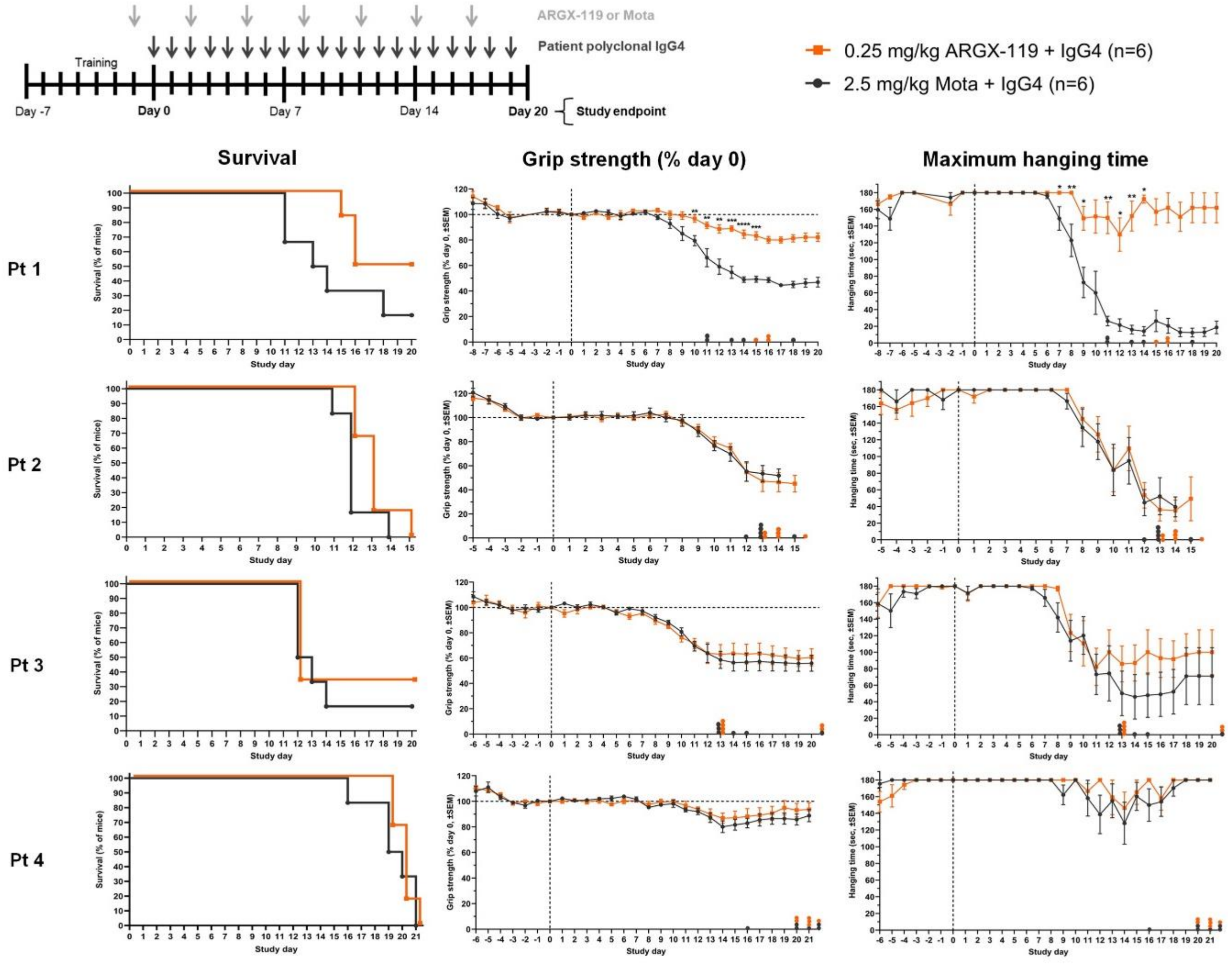


**Supplementary Figure 4: ARGX-119 rescues the MuSK MG phenotype induced with polyclonal patient IgG4 in a patient-specific manner.** *From left to right for patients 1-4*: percentage mouse survival over time, with reductions in survival representing sacrifice of mice due to humane endpoint reached; percentage grip strength normalized to day 0; and hanging time (maximum 180 seconds, best of 3 attempts) in mice treated with 0.25 mg/kg ARGX-119 or isotype control Mota. Data points of percentage grip strength and hanging time represent grouped values of the mean ±SEM of mice. Data points on the x-axis represent mice that were sacrificed as they reached humane endpoint or end of study. Statistical analyses were only performed on average data points when the groups contained more than three mice. *P<0.05, **P<0.01, ***P<0.001, ****P<0.0001 for ARGX-119 vs Mota treated mice.

|  | Mouse Fz-domain |  | Human Fz-domain |  |
|---|---|---|---|---|
| Material | Binding of IgG4 alone (RU) | % competition with ARGX-119 | Binding of IgG4 alone (RU) | % competition with ARGX-119 |
| Healthy donor IgG4 | 0 | 0 | 0 | 0 |
| Pt1 IgG4 | 4 | 0 | 10 | 0 |
| Pt 2 IgG4 | 33 | 0 | 109 | 8.3 |
| Pt 3 IgG4 | 0 | 0 | 2 | 0 |
| Pt 4 IgG4 | 0 | 0 | 10 | 0 |

**Supplementary Figure 5: SPR data quantifying competition of ARGX-119 and patient IgG4 for MuSK-Fz domain binding.**

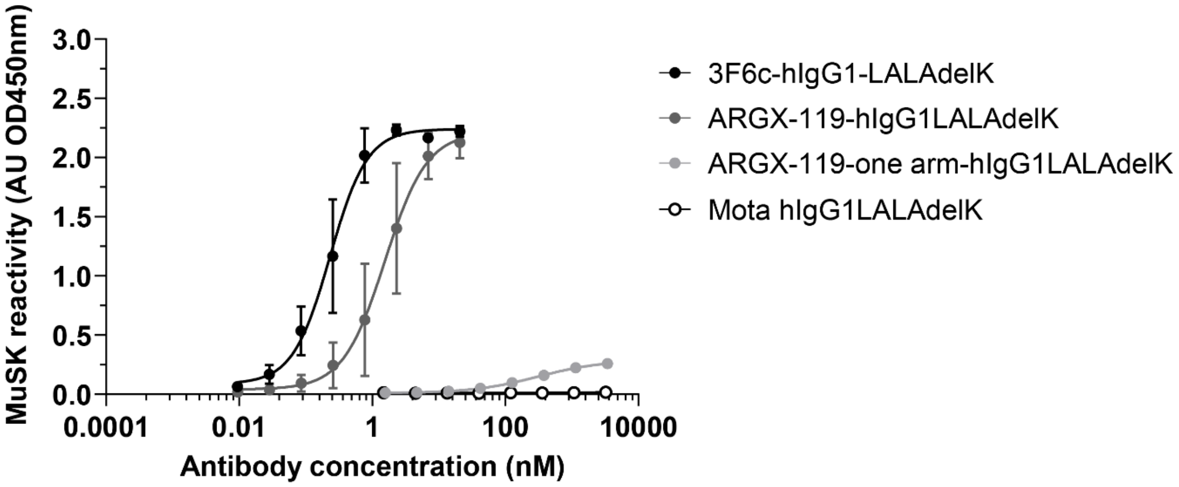


**Supplementary Figure 6: Confirmation of the MuSK bio-bridging assay design.** Bivalent MuSK antibodies 3F6c-hIgG1-LALAdelK and ARGX-119 hIgG1LALAdelK serve as positive controls in this bio-bridging assay, whereas a one-armed ARGX-119 antibody and the isotype control antibody Mota show no to little binding in this assay.

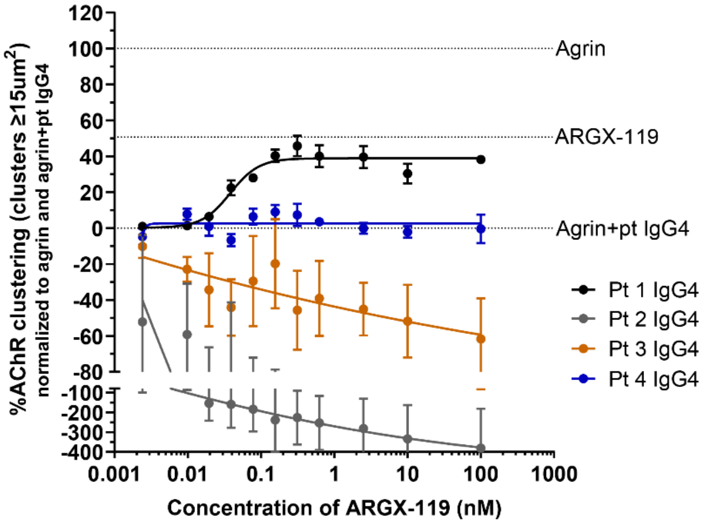


**Supplementary Figure 7: Dose-response curve of ARGX-119 in an *in vitro* MuSK MG model.**

Dose-response curve of ARGX-119 on AChR clustering (clusters ≥15um$^2$) in C2C12 myotubes treated with pt IgG4s. The dashed lines represent conditions where agrin or ARGX-119 were added to the cultures alone. The data was related to the inhibitory capacity of each of the patient IgG4s individually. Data represent mean ± SEM.
